# Temporal effects on the abundance of lethal fungal pathogens of amphibians

**DOI:** 10.1101/2025.08.08.669422

**Authors:** Sarah E. Dingel, Samina F. Hanif, Scott W. Greenhalgh

## Abstract

Fungal pathogens greatly threaten amphibian diversity, bringing many species near extinction. Consequently, informing on the abundance and ecological characteristics of fungal pathogens, such as Batrachochytrium dendrobatidis (Bd), through tools such as mathematical models is of the utmost importance with respect to current conservation efforts. While traditional mathematical models, such as the logistic growth model, can infer basic details on Bd abundance, the model’s simplistic nature renders it unable to account for myriad external factors. So, to inform on some of these factors, namely temporal fluctuations in the Bd growth rate, and carrying capacity, we extended the logistic growth model to consider combinations of time-varying coefficients. For our new models, we estimated model parameters from publicly available data on Bd zoospore density across multiple temperature ranges and geographies, assessing the quality of model fit relative to complexity by Akaike Information Criterion, and Akaike weights, in addition to characterizing potential long-term behaviors through stability analysis. Our work shows that our time-varying growth rate and carrying capacity model was at least 1.4 times more likely to reflect Bd abundance at optimal temperature ranges. This suggests a multi-pronged approach for hindering Bd, namely at non-optimal temperatures, conservation efforts such as tadpole removal and water disinfection should be utilized consistently, and at optimal temperatures, they should be timed to when they generate the greatest benefit with respect to the elimination of Bd zoospores.

## 1. Introduction

Imagine a world without amphibians. Such loss in biodiversity would irreparably damage the ecological food chain, likely causing mass extinction events for a myriad of other species [1]. While this scenario may seem far-fetched, lethal pathogens exist that endanger over 43% of amphibian species worldwide [2]. For instance, one such pathogen is *Batrachochytrium dendrobatidis* (Bd), which causes Chytridiomycosis in the amphibian species it infects. To elaborate, when Bd infects amphibians, they lose the ability to regulate their salt content and utilize their skin as a breathing mechanism, ultimately leading to death [3].

The life cycle of Bd consists of two stages: a substrate-dependent zoosporangium stage and a zoospore stage. Only zoospores infect amphibians, which do so by attacking the keratinized tissue in their skin cells where they mature into zoosporangia. The zoosporangia either propagate into the environment to infect other organisms [4], or reinfect the original host, exasperating infection intensity [5].

The capacity for Bd to infect amphibians depends on various factors [6], with recent work illustrating the role that temperature and geography play in Bd zoospore abundance [4]. Specifically, Bd abundance estimates were shown to vary across Louisiana (LA), New Mexico (NM), Ohio (OH), Tennessee (TN), and Vermont (VT), and at temperatures of 4, 12, 17, 21, 25, 26, and 27 degrees Celsius, respectively, by quantifying zoospore density through MTT assays [4]. In brief, an MTT assay is a colorimetric assay that assesses the number of healthy cells, or under the context of Bd, zoospores, in a sample. For each sample, active cells reduce MTT salt into Formazan, causing a color change from yellow to purple, which can then be quantified by a spectrophotometer [7], and ultimately serve as a proxy for zoospore density.

One approach to describe the evolution of Bd zoospore density is to utilize the logistic growth model [4]. The logistic growth model is ubiquitous in modeling biological phenomena [8], as it is often used first to forecast the population growth of bacteria, fungi, and viruses [8]. Although the logistic growth model provides a solid baseline for such work, it does not take into account external factors that may impact reproduction and abundance, such as temperature, and humidity [6], among others [9]. Consequently, more complex models are often proposed to describe the impacts of external factors, typically at the cost of requiring specialist mathematical theory to properly analyze. Fortunately, not all generalizations require specialization, as the simple extension of the logistic growth model to one that considers combinations of a time-varying intrinsic growth rate and carrying capacity can often be analyzed using standard techniques. One of the most recognized uses of the logistic growth model within epidemiology is the modeling of fungal spread [10]. With a betterfit model, we are able to more accurately model and predict the spread of Bd. The density of viable zoospores of Bd is impacted by the surrounding temperatures, whether it fosters growth or inhibits spread. Our model allows a better showcase of this fluctuation in the growth rate and carrying capacity, as we account for the temporality of the environment, when compared to the traditional logistic growth model at temperatures higher than 4°C.

The growth rate and carrying capacity of a population is subject to various exogenous elements. For example, biological mechanisms often have ideal environments and temperatures for metabolic activities [11]. Additionally, environmental fluctuations can reflect the ability to sustain a certain amount of life and contribute to variations in the carrying capacity, given the resources available. Since the elements of growth rate and carrying capacity are biological phenomena, it is fair to assume periodic fluctuations in response to temporality [8].

While prior work has assessed thermal and geographic effects on zoospore density utilizing logistic growth models [4], it does not take into account external factors that may influence Bd reproduction or abundance. However, it is well known that external factors may cause temporal variations in such characteristics [6]. To inform on this issue, we modified the logistic growth model to include combinations of a time-varying intrinsic growth rate and carrying capacity.

With these new models, we examine the quality of fit to Bd zoospore data relative to model complexity, to inform on what characteristics play an integral role in Bd abundance, and thereby contribute to the information needed by ecologists to make informed decisions for protecting amphibian populations.

## 2. Methods

To model the temporal effects of Bd abundance, we extended the original logistic growth model to include time-varying coefficients. Specifically, we consider scenarios where either the growth rate or the carrying capacity varies by time and a scenario where both variables are time-varying. We calibrate each scenario to data on the Bd zoospore viability [4], specifically in regions where the primary host species was of the *Lithobates* genus. To determine the optimal model relative to complexity, models were compared using AIC scores, with the logistic growth model AIC value serving as the baseline. Modeling scenarios and analysis were implemented into R.

### 2.1 Logistic growth model

For the purposes of all the models, x(t) represents the population density of the zoospores. The logistic growth model is defined as

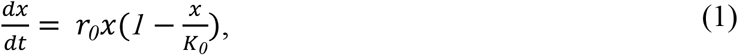

where r_0_ represents the intrinsic growth rate of the zoospore, *K*_*0*_ represents the carrying capacity with respect to environmental conditions, and *x* is the zoospore density.

### 2.2 Time-varying growth rate model

We first generalized the logistic growth model to a time-varying growth rate (TVGR) model. Specifically, we assumed it varies temporally and is defined by

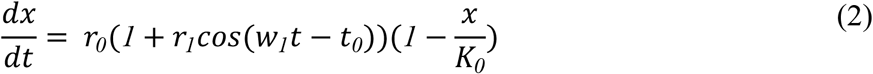

where *r*_*0*_ is the intrinsic growth rate, *r*_*1*_is the amplitude of temporal effects on the growth rate, and w is the frequency of temporal effects.

### 2.3 Time-varying carrying capacity model

Our next generalization considers a time-varying carrying capacity (TVCC) model, as given by,

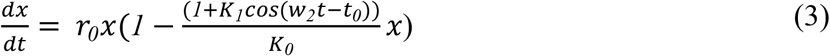

Here, *K*_*0*_ denotes the carrying capacity without temporal effects, *K*_*1*_ represents the amplitude of temporal effects on the carrying capacity, and w is the frequency of temporal effects.

### 2.4 Time-varying growth rate and time-varying carrying capacity model

The full model combines both the time-varying growth rate and time-varying carrying capacity (TVGRCC),

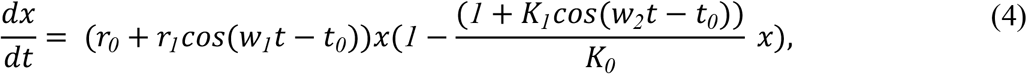

where the parameters are as defined in sections 2.1.2 and 2.1.3.

### 2.5 Estimation of parameters

To determine parameters we fit each model to datasets, utilizing least squares minimization [12]. Specifically, for each model, the optimal parameters were identified by minimizing the error of its predictions relative to data,

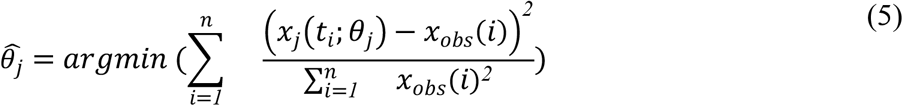

where 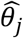 is the optimal parameters for the *j*^*th*^ model, *x*_*obs*_(*i*) is the observed MTT zoospore density at time *i* from a particular isolate at a given temperature, and *x*_*j*_(*t*_*i*_; *θ*_*j*_) is the estimated zoospore density generated by the *j*^*th*^ model, given the parameter set *θ*_*j*_.

Optimal parameters were estimated using R’s optim() function (Figure 2, Figure 2, Supplemental materials). For estimating optimal parameters and initial conditions of the logistic growth model (1), parameter ranges were based on biologically feasible values from data and the literature. Subsequent models utilized similar procedures for the TVGR, TVCC, and TVGRCC models, with initial parameter estimates corresponding to the optimal values from the fit of the logistic growth model (1) as initial parameter estimates.

**Figure 1).**
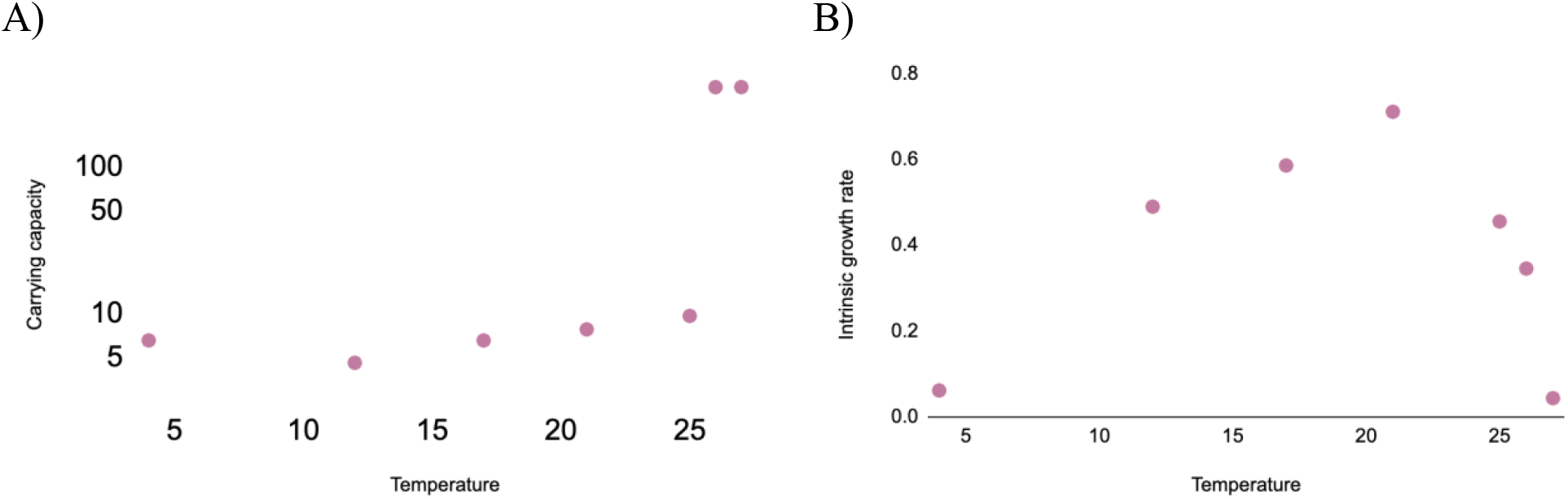
LA *r*_*0*_and *K*_*0*_ estimates. The estimates of A) average carrying capacity, and B) average intrinsic growth rate over temperature (red dots). Estimates for the NM, OH, and TN isolates can be found in the supplementary materials.

**Figure 2).**
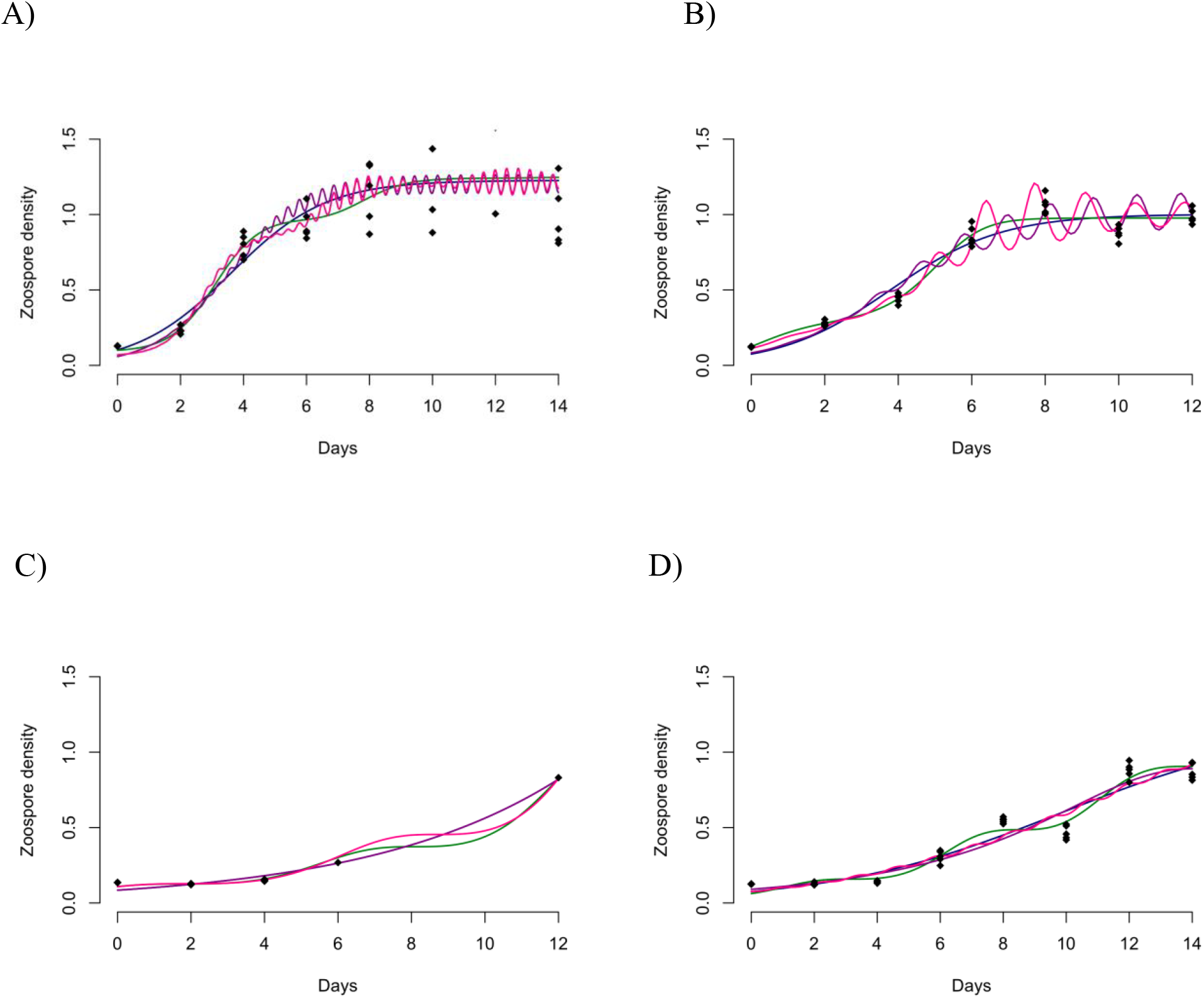
Trajectories of zoospore density. The trajectories of zoospore density over time for A) LA, B) NM, C) OH, and D) TN isolates at 21°C, for the logistic model (blue curve), TVGR (green curve), TVCC (purple curve), TVGRCC (pink curve), and collected zoospore data (black dots). The trajectories for the isolates at the remaining six temperatures can be found in the supplementary materials.

### 2.6 Akaike information criteria

To assess the quality of each model’s fit to data we calculate AIC values and scores. AIC scores are a measure that evaluates the fit of a model relative to the number of parameters used [13]. Defining the sums of squares error (*SSE)* for the *j*^*th*^model,

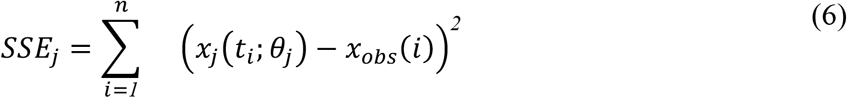

it follows that of the AIC value for the *j*^th^ model is

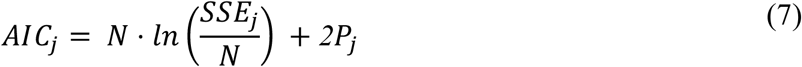

where *P*_*j*_ is the number of parameters, and *N* is the number of data points.

To obtain the AIC score for each model, we subtract AIC values from the logistic growth model’s AIC value for each particular dataset,

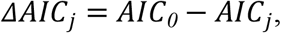

where *AIC*_*0*_ is the AIC value of the logistic growth model.

We calculate AIC scores for each isolate and temperature, across all isolates for a given temperature, across all temperatures for a given isolate, and across all isolates and temperatures.

In addition to AIC scores, we calculate the Akaike weights of the four models. AIC weights provide an approximation of the probability that a model is best given a set of candidate models [14]. An Akaike weight for the *i*^*th*^ model is defined as,

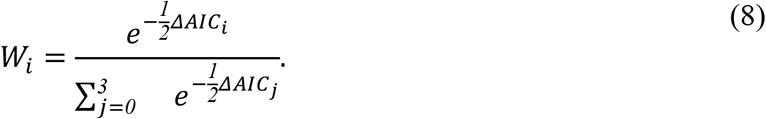

### 2.7 Stability analysis

To classify potential long-term behaviors of models, we perform stability analysis [15]. For models with a closed-form solution, we provided these conditions simply by examining the potential limiting behaviors as time approaches infinity.

For models without closed-form solutions, namely the TVGRCC model, we provided a Poincare map to characterize its long-term solution behaviors [16]. Specifically, the Poincare map of the TVGRCC model is of the form

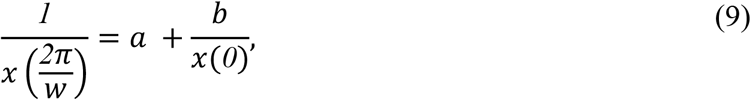

where the constants *a* and b are uniquely determined from the time-varying coefficients of the TVGRCC model [15].

## 3. Results

We assessed the impacts of temporality on the growth of Bd across four isolates at seven different temperatures by introducing three mathematical logistic growth models. These models were aligned to publicly available data on zoospore density and applied to help estimate AIC values, stability, and inform on temporal impacts on Bd growth.

### 3.1 Model solutions and stability properties

Below we characterize conditions for all long term behavior of our four models. We start by presenting the criteria for models with closed-form solutions, namely the logistic, TVGR, and TVCC models, followed by the TVGRCC model through the analysis of its Poincare map (Table 1).

**Table 1.**
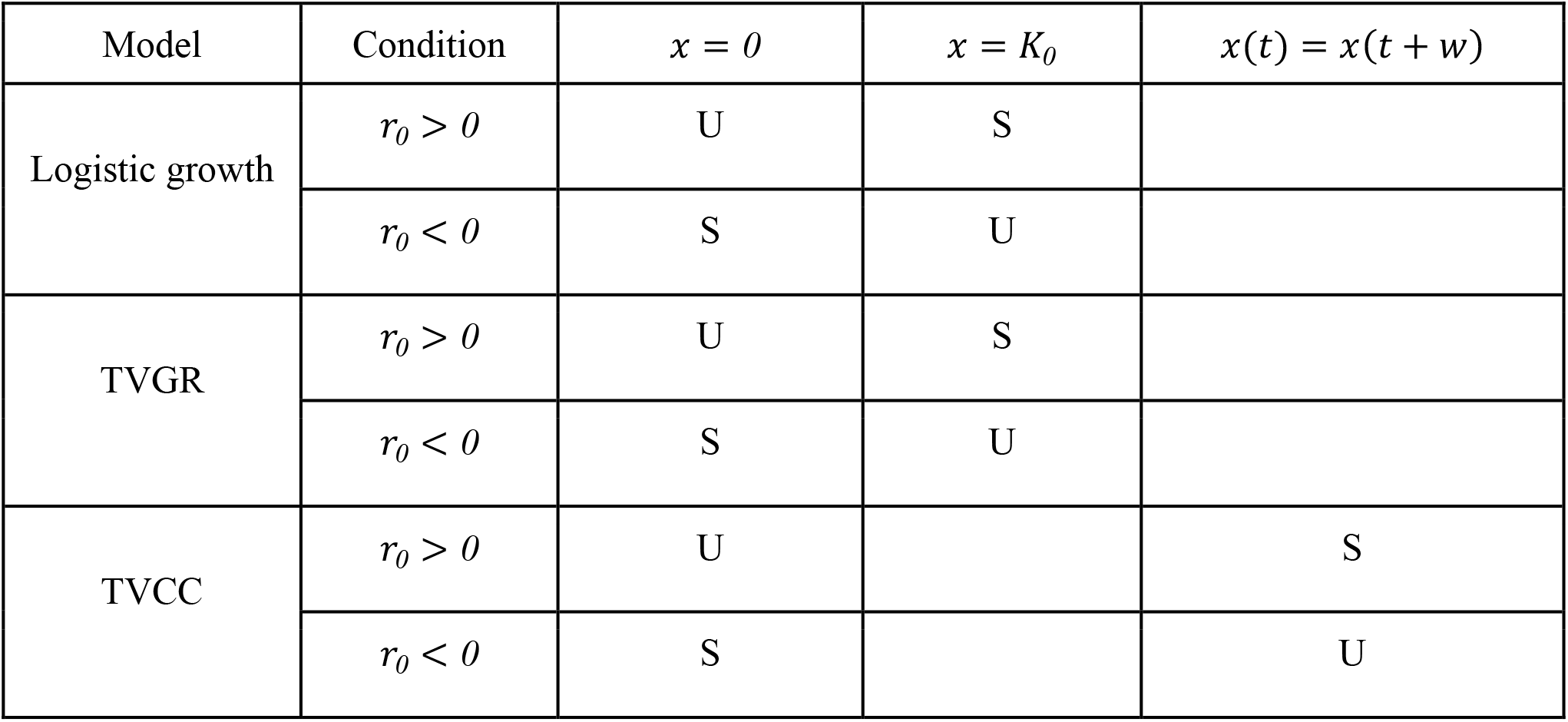
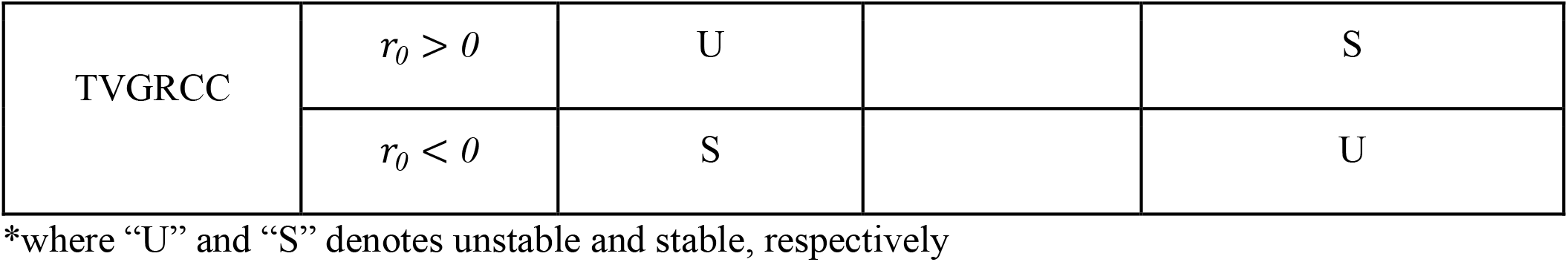
Long-term solution behavior for each model.

| Model | Condition | $x = 0$ | $x = K_0$ | $x(t) = x(t + w)$ |
| --- | --- | --- | --- | --- |
| Logistic growth | $r_0 > 0$ | U | S | |
| | $r_0 < 0$ | S | U | |
| TVGR | $r_0 > 0$ | U | S | |
| | $r_0 < 0$ | S | U | |
| TVCC | $r_0 > 0$ | U | | S |
| | $r_0 < 0$ | S | | U |
| TVGRCC | $r_0 > 0$ | U | | S |
| | $r_0 < 0$ | S | | U |

\*where “U” and “S” denotes unstable and stable, respectively

#### 3.1.1 Long-term behavior of models with closed-form solutions

The solution of the logistic growth model is

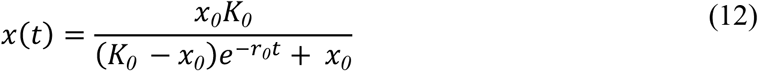

Taking the limit as *t* → *∞*, it follows for *r*_*0*_ > *0* that *x*(*t*) → *K*_*0*_ and for *r*_*0*_ < *0* that *x*(*t*) → *0*.

Turning to the TVGR model, it has the closed-form solution,

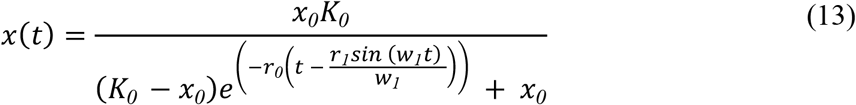

For *r*_*0*_ < *0*, we have that *x*(*t*) → *0* and when *r*_*0*_ > *0* it follows that *x*(*t*) → *K*_*0*_ as *t* → *∞*. The TVCC model has the solution,

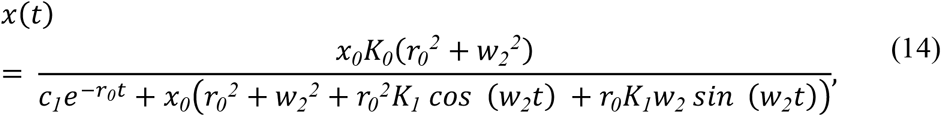

where we define 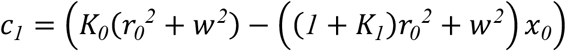 for the ease of presentation. As *t* → *∞, x*(*t*) → *0* for *r*_*0*_ < *0*, and for *r*_*0*_ > *0*, we have that *x*(*t*) approaches the periodic solution

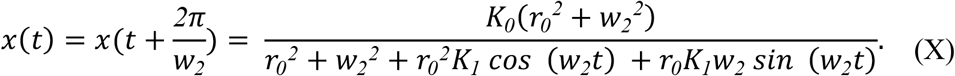

#### 3.1.2 Long-term behavior of the TVGRCC model

The TVGRCC model does not possess a closed-form solution. To analyze its long term behavior, we consider its Poincare map (9), reformulated as the recurrence equation

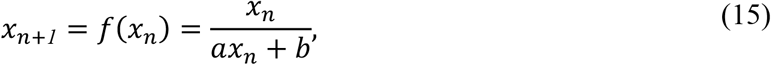

Defining α(*t*) = *r*_*0*_ (*1* + *r*_*1*_ *cos*(*w*_*1*_ *t* − *t*_*0*_)) and 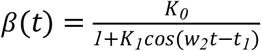, we have the formula for *a*is

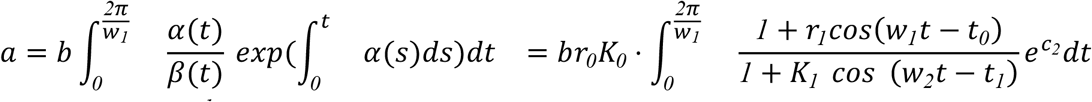

Where 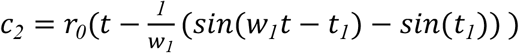, and the formula for *b* is

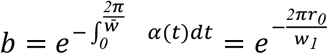

where *n* and *m* are the smallest integers *s*. *t*. *<u>w</u>* = *nw*_*1*_ = *mw*_*2*_ .

The recurrence Eq (15) has two equilibrium points, 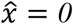, and 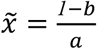.

The stability of Eq (15) near 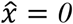, follows from

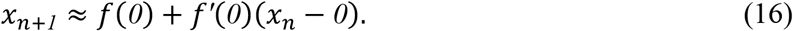

Thus, provided 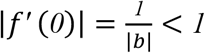, or equivalently |*b*| > *1*, we have that 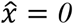 is stable, and *x*(*t*) → *0*.

To inform on the stability of Eq (15) about 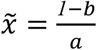, we have that

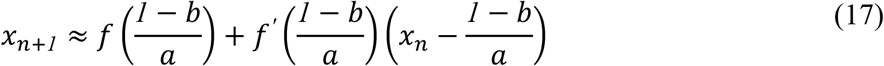

Provided 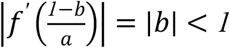 we have that 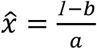 is stable.

Note that |*b*| < *1* implies the TVGRCC model approaches a stable periodic solution when *r*_*0*_ > *0* as *t* → *∞*, and |*b*| > *1* implies that the extinction equilibrium of *x* = *0* is stable for values of *r*_*0*_ < *0*.

### 3.2 Akaike information criterion

#### 3.2.1 AIC scores for specific isolates and temperatures

No detectable pattern was found alluding to the idea that one model is better than another, across all isolates and temperature, with the exception of Eq (1) at a temperature of 4°C, with AIC scores of -462.3636 for LA, -619.2097 for NM, -696.6526 for OH, and -775.0110 for TN (Supplementary materials).

#### 3.2.22 AIC scores across all isolates

When comparing AIC scores across isolates, the smallest AIC scores were -1665.4215, - 770.5017, and -718.6932 for Eq (4) at temperatures of 12, 17, and 21, respectively. Similarly, the lowest scores for temperatures of 25, 26, and 27 were -854.2198, -808.8926, and -1031.6234, respectively and correspond to Eq (2) (Table 2).

**Table 2.**
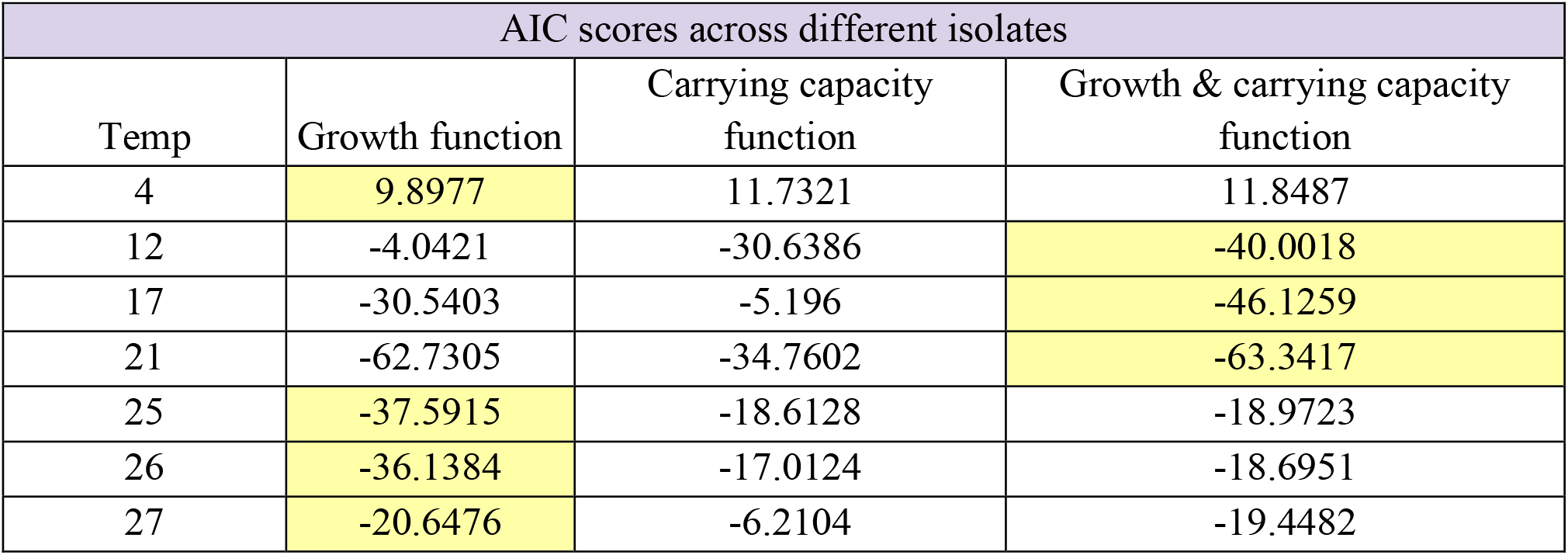
AIC score across isolate.

#### 3.2.3 All temperatures

When classifying by isolate, Eq (4) had the lowest AIC scores for LA, NM, and OH, with respective values of -1720.834, -2079.534, and -2279.569. TN was the only isolate that had the lowest AIC value aligned with Eq (2) with a value of -2335.728 as seen in Table 3.

**Table 3.**
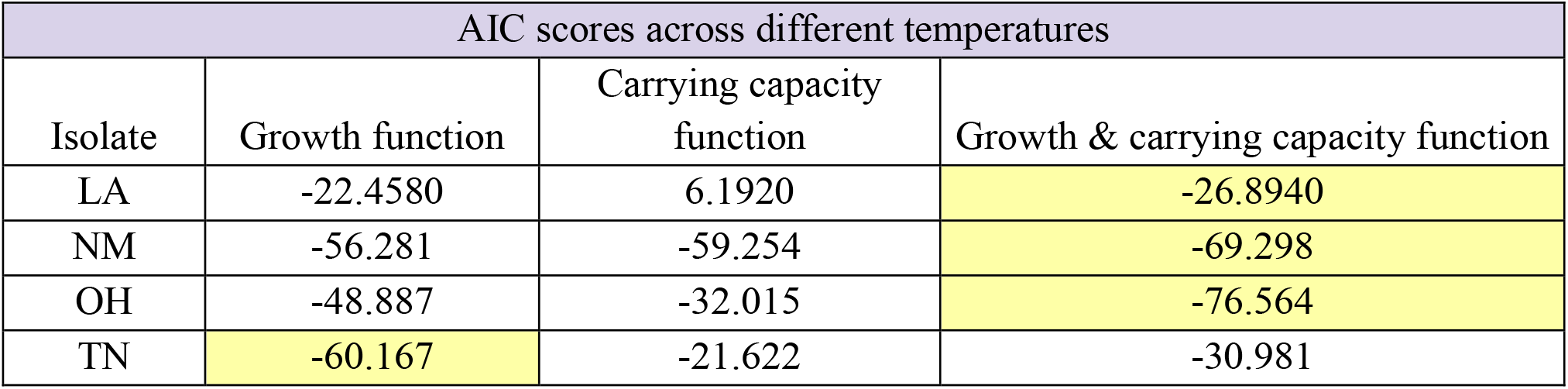
AIC score across temperature.

#### 3.2.4 AIC scores across all isolates and temperatures

#### 3.2.5 Akaike weights

Using Akaike weights, it was generally seen across the four isolates that the TVGRCC had the highest probability of being the best fit, namely with OH, NM, and LA, represented in Figure 3A. TN is the outlier, as its best fitting model is the TVGR. Across temperature, there was more variability, where both TVGR and TVGRCC were largely acceptable models, as shown in Figure 3B. TVGR was the best model for temperatures 12, 17, and 21°C, while the TVGRCC was the best model at temperatures of 25, 26, and 27 °C. At 4 °C, graphed across isolates, there is a low probability that any of the models are good fits, with a probability of fit of ∼0.007, attributed to the TVGR model.

**Figure 3).**
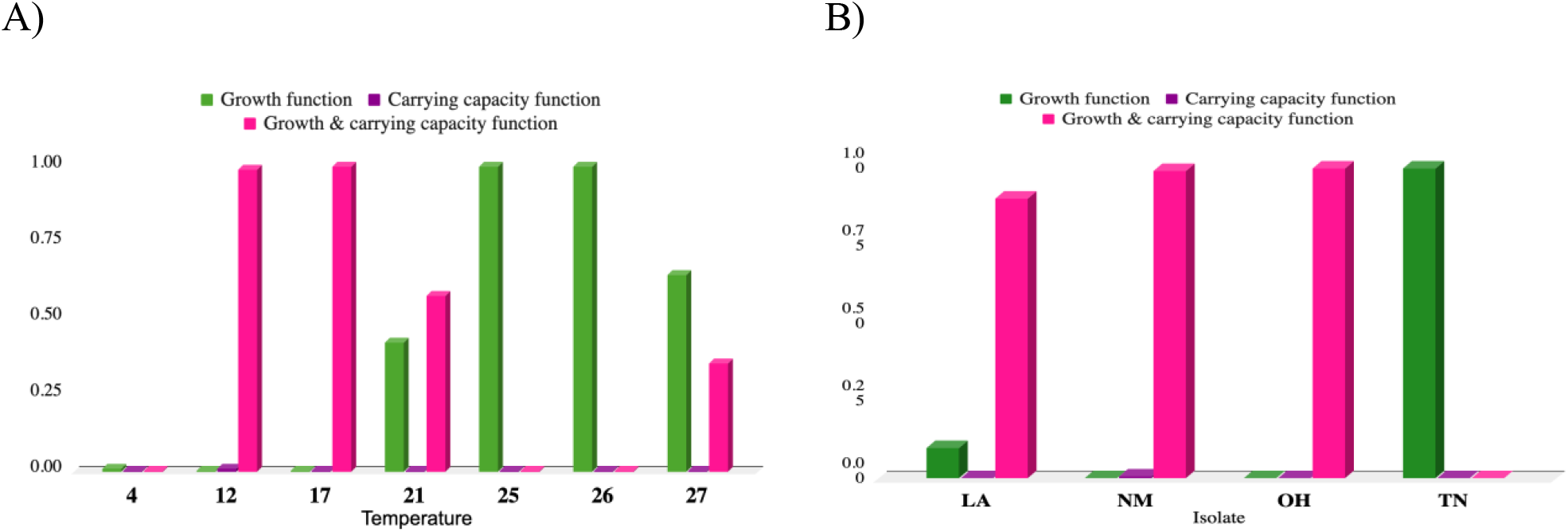
Akaike weights across isolates and temperatures. The probability a model is the best fit A) across each temperature and B) across LA, NM, OH, and TN isolates.

### 3.3 Parameter estimates

Bd zoospore data collected from NM, OH, and TN all showed optimal intrinsic growth around 25 °C and an optimal carrying capacity around 5°C (see supplementary materials). LA, on the other hand, showed optimal growth at a range of 15-20 °C and optimal carrying capacity conditions between 25-27 °C (Figure 3).

## 4. Discussion

We evaluated four mathematical models to demonstrate the impact of temporality of the population characteristics of Bd zoospores by different temperature conditions and geographies. For our models, we characterized the stability of their long term behaviors, and utilized AIC scores and Akaike weights to inform on which model was best suited for its application on fungal growth. We illustrated logistic growth models with closed-form solutions extending beyond the generic logistic growth model for the TVGR and TVCC model, using limits to analyze its stability. For the TVGRCC model, there was no closed-form solution and its stability was determined by the use of Poincare maps.

With recent declines in the amphibian population, understanding the characteristics of Bd and how to mitigate its spread is important in protecting at-risk species. Our work informs on this concern by the addition of periodic parameters in model dynamics, which allows for better understanding of the impacts of temporality on intrinsic growth rate and carrying capacity. While it is uncertain what the exact temporal rates are for Bd’s intrinsic growth rate and carrying capacity, our periodic models generally fit the data on zoospore density better than the logistic growth model, even with complexity considerations, according to AIC scores, which eluded to time-varying models being more complex and therefore being a better fit.

There are many circumstances that can impact predictions of Bd at all isolates and all temperatures. For instance, it is known that loss of habitat that could result from droughts or urbanization can affect the growth rate or the carrying capacity of populations. AIC scores showed that not one model trumped another, which is to be expected with such considerations. For a time-varying growth rate, this would indicate that temporal shifts are leading to a change in the rate at which Bd are able to reproduce and grow that could have further biological implications due to the activity of their metabolic enzymes at particular temperatures. Often, different organisms possess enzymes with alternate optimal conditions for carrying out their processes. Enzymes are proteins, which are moderately sensitive to temperature, pH, and other variables, as they have a range of functionality. Once the conditions venture out of that range, they tend to have a sharp decline in their productivity [2]. Considering that our isolates varied across the US, we can assume that the Bd growth rate at each isolate will vary.

Certain environmental conditions can contribute to an increase or decrease of the intrinsic growth rate and carrying capacity. At optimal temperatures, Bd’s growth rate and carrying capacity efficiently propagate spread. Conversely, at suboptimal temperatures, occurring naturally or due to human intervention, the ability of Bd to procreate and infect more amphibians is likely to decrease. Our results corroborate this when at the suboptimal temperature of 4°C, the logistic growth model performed the best in fitting the data. While the logistic model performs relatively well at suboptimal temperatures, caution should be taken as fungi possess the ability to evolve and undergo natural selection to conform to their new environmental temperatures [9].

The similarity of Bd to other fungal pathogens, such as *Batrachochytrium salamandrivorans* (Bsal), implies they may follow similar temporal patterns across the geographies and temperatures that we studied. Furthermore, many fungal pathogens share similar infection capabilities due to certain molecular continuities. For example, heat shock proteins may aid in Bd and Bsal’s ability to resist cessation when external temperatures rise.

While we considered simple cosine curves in our time-varying rates, it could be generalized to many other time-varying forms. For example, we could generalize the models based on the Fourier series in order to better capture the time-varying behavior of the model [19]. Another extension could be the Lotka-Volterra model [20], which would allow for the consideration of other external factors and inter species interaction. This would potentially allow for a better biological inference. We could also consider a factorial experiment, as an additional form of analysis, providing deeper insight into uncertainty and sensitivity analysis.

The LA isolate was taken from the host species of *Acris crepitans*, while the other four isolates were taken from a variety of different species of the *Lithobates* genus [4]. LA was considered an outlier, as the estimates of its parameter values were much larger than the others. This difference in host species could account for the extensively large *K*_*0*_ values that were seen for temperatures 25, 26, and 27 °C, which rendered the graphs and models inaccurate.

As with all models, our model involves certain simplifications. For instance, our model is a closed system and does not permit extreme environmental changes or black swan events. To elaborate, any natural disaster would wipe out host species, fungal pathogens, and many other factors that are important in their ecology, which are not accounted for in our models’ growth rate or carrying capacity. Specifically, our rates can not compensate for such abrupt changes to the environment, without the inclusion of impulse effects. Moreover, our models’ may not accurately represent temperature variability over a certain threshold, as the data utilized was over a temperature range of 4-27 °C. Caution should also be taken with extending the work to other Bd strains, as data in our analysis was only collected from two Bd strains within its 6 known phylogenetic lineages [4,21]. Another weakness of our data was the MTT assay that was utilized as a proxy for zoospore density. Under the assumption that each zoospore is not limited to one single mitochondria, there is no known ratio between viability and absorbance, as we lack sufficient information on Bd mitochondrial activity and abundance. However, despite these various limitations, with modifications to the rates, our models’ could be fashioned to provide insight on many of these issues with modest efforts.

In summary, understanding Bd will allow for the development of better conservation efforts to mitigate its impact on amphibians. Our work puts forward a multi-pronged approach for constraining the abundance of Bd. Specifically, at non-ideal temperatures, preservation efforts, including tadpole removal and water disinfection should be utilized consistently, while at ideal temperatures, efforts should be timed to when they will yield the greatest benefit with respect to the eradication of Bd zoospores.

## Supporting information

Supplemental methods and results

## Use of AI tools declaration

The Authors declare they have used Grammarly, as a form of artificial intelligence.

## Acknowledgements

We would like to thank…

National Science Foundation, Dr. Clark, Siena College

## Conflict of interest

The authors declare there is no conflict of interest

## Notes

### Competing Interest Statement

The authors have declared no competing interest.

