## Supplemental methods and results for "Temporal effects on the abundance of lethal fungal pathogens of amphibians"

### Supplementary materials

The following document provides additional information on the fitting of the mathematical models and AIC estimations

#### Isolate graphs across all temperatures

All four mathematical models plotted against the collected zoospore data are shown below. Models are differentiated by color to indicate which model has the closest fit to the data.The blue line represents the basic logistic model, the green line shows the time-varying growth rate model, purple shows the time-varying carrying capacity model, and lastly, the pink line indicates the time-varying growth and carrying capacity model.

LA 4

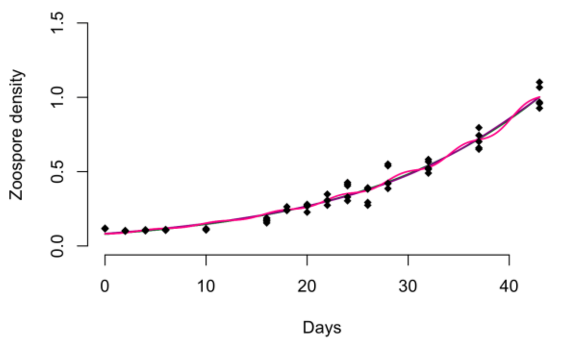


LA 12

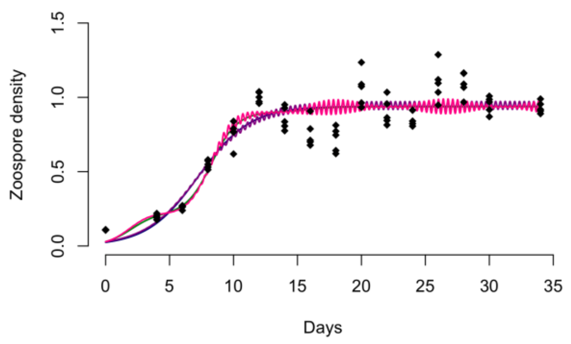


LA 17

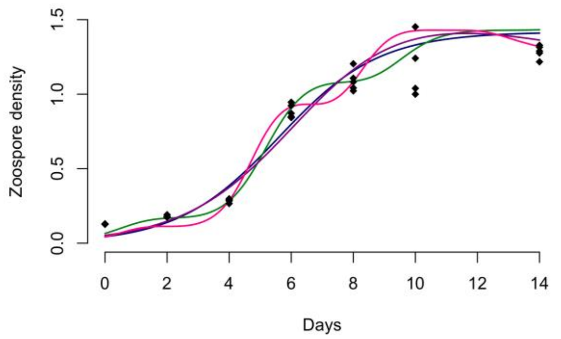


LA 21

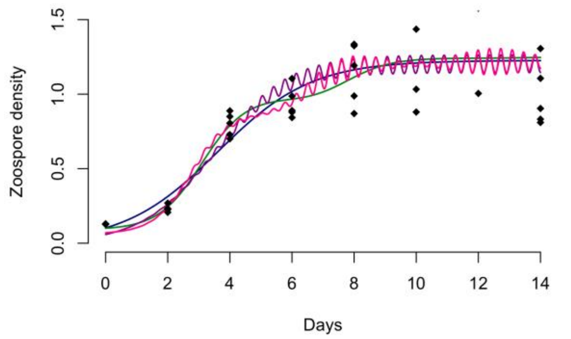

LA 25

**inaccurate graph as the parameter estimates were much larger than the other isolates

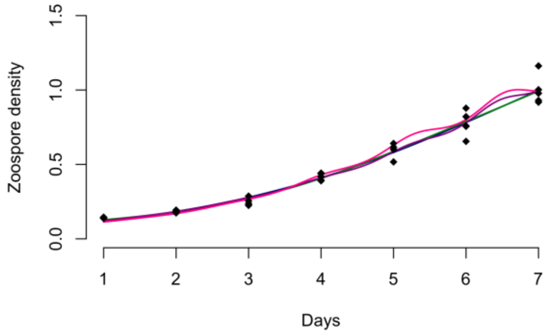


LA 26

**inaccurate graph as the parameter estimates were much larger than the other isolates

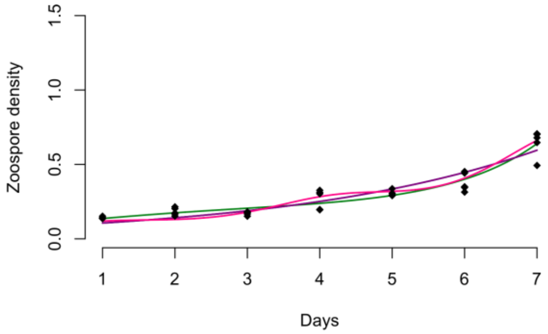


LA 27

**inaccurate graph as the parameter estimates were much larger than the other isolates

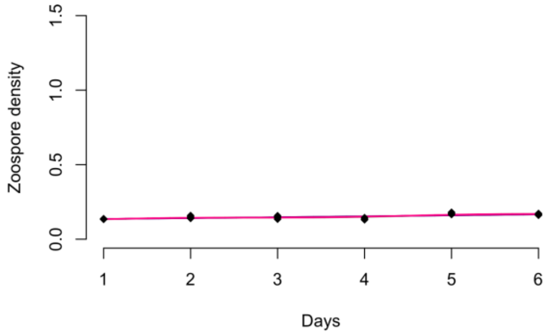

NM 4

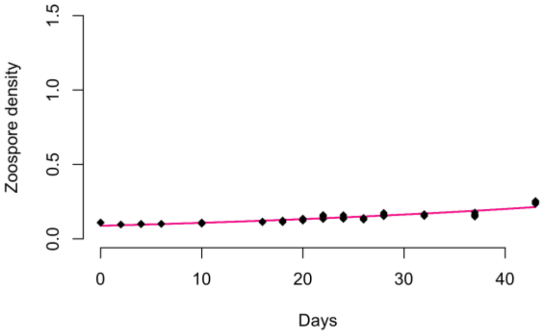


NM 12

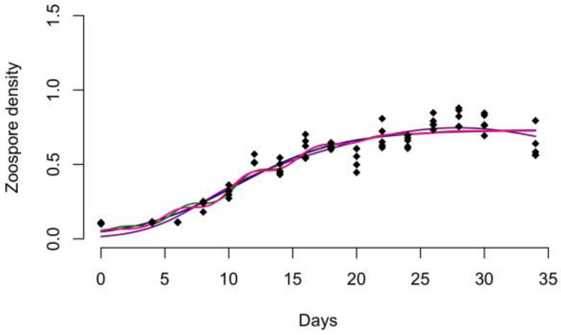


NM 17

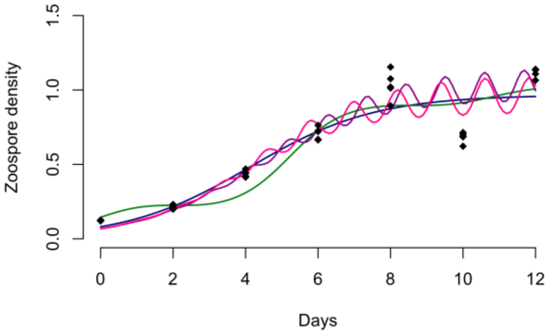


NM 21

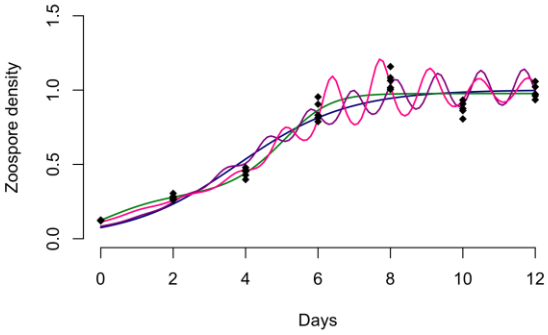


NM 25

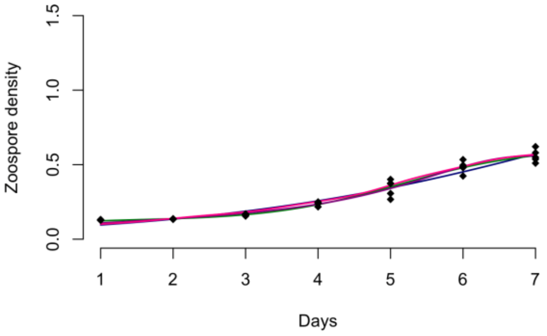


NM 26

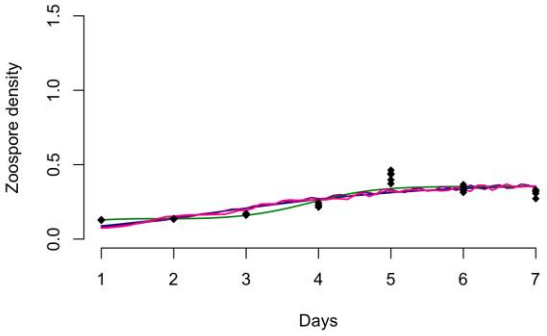

NM 27

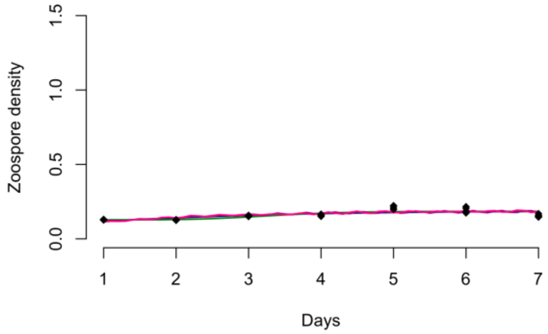

OH 4

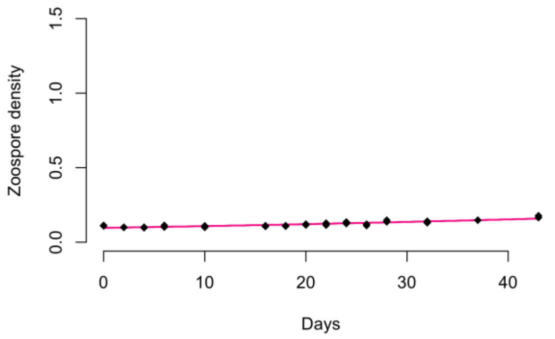


OH 12

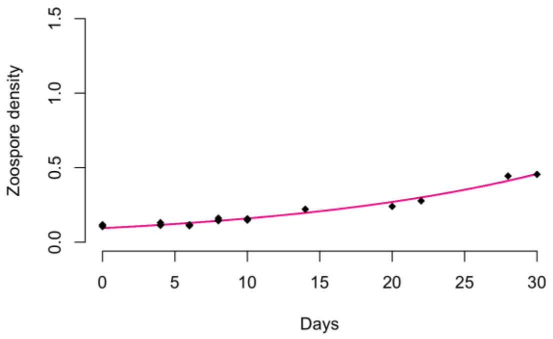


OH 17

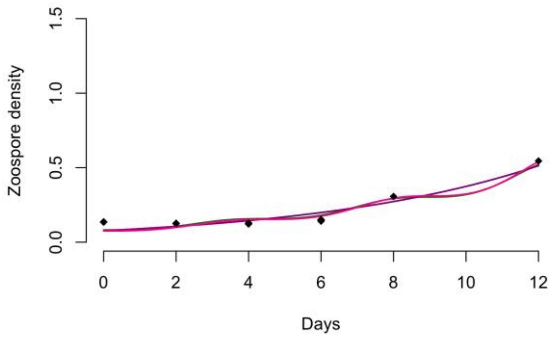


OH 21

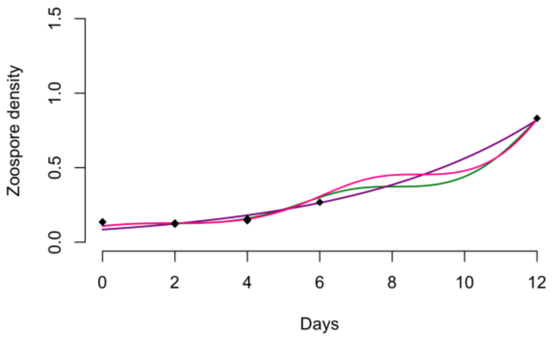


OH 25

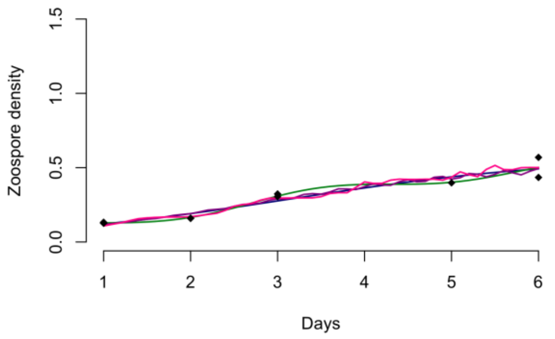


OH 26

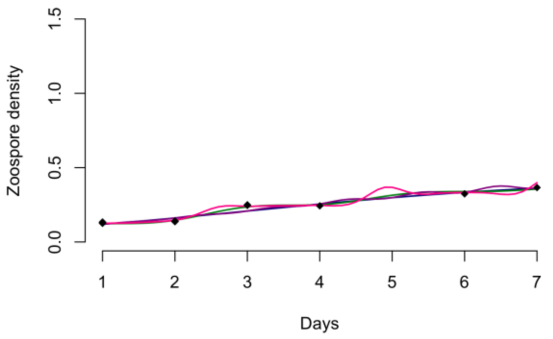

OH 27

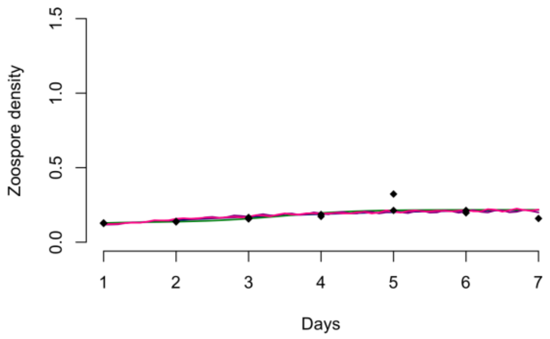

TN 4

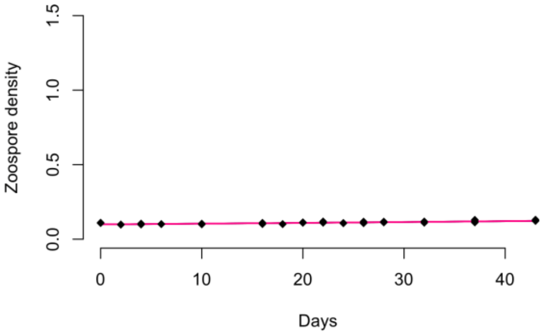


TN 12

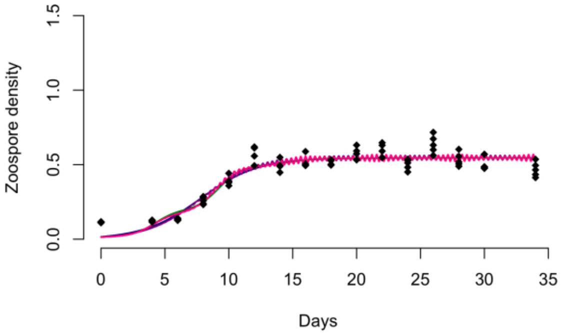


TN 17

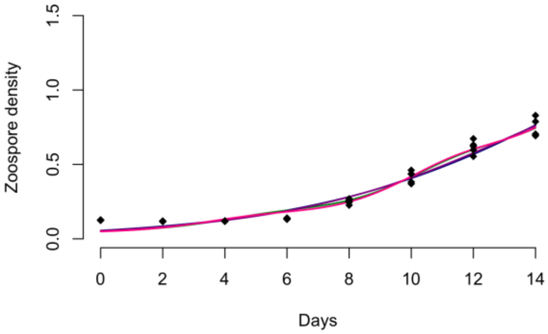


TN 21

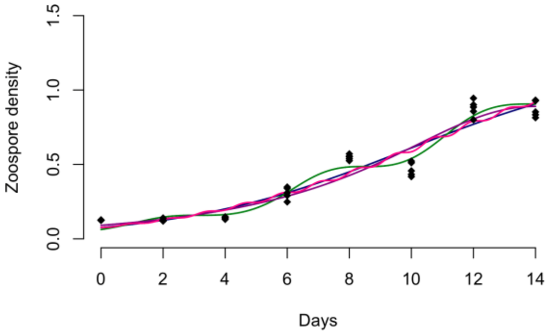


TN 25

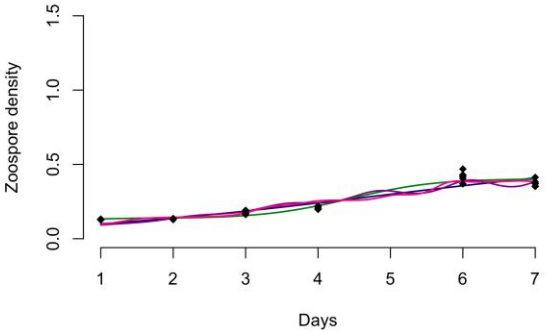


TN 26

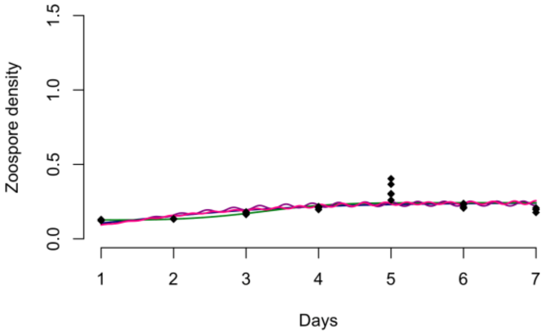


TN 27

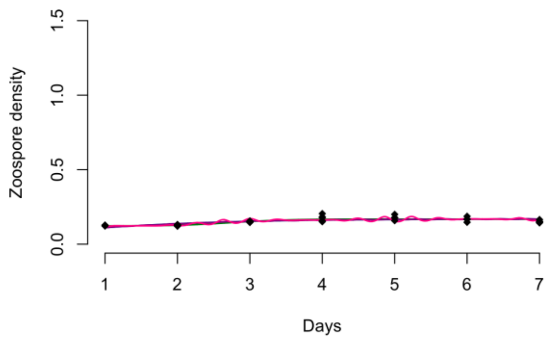

1.5
1.0
0.5
0.0
Zoospore density
1
2
3
4
5
6
7
Days

### Estimating $r_0$ and $K_0$ parameter values for isolates

To model trends in growth rate and carrying, parameter values were graphed against temperature to speculate on optimal conditions for zoospore growth and sustainability. LA and TN are part of the GPL1 strain of Bd, while NM and OH are part of the GPL2 strain. Exact values can be found in the R script.

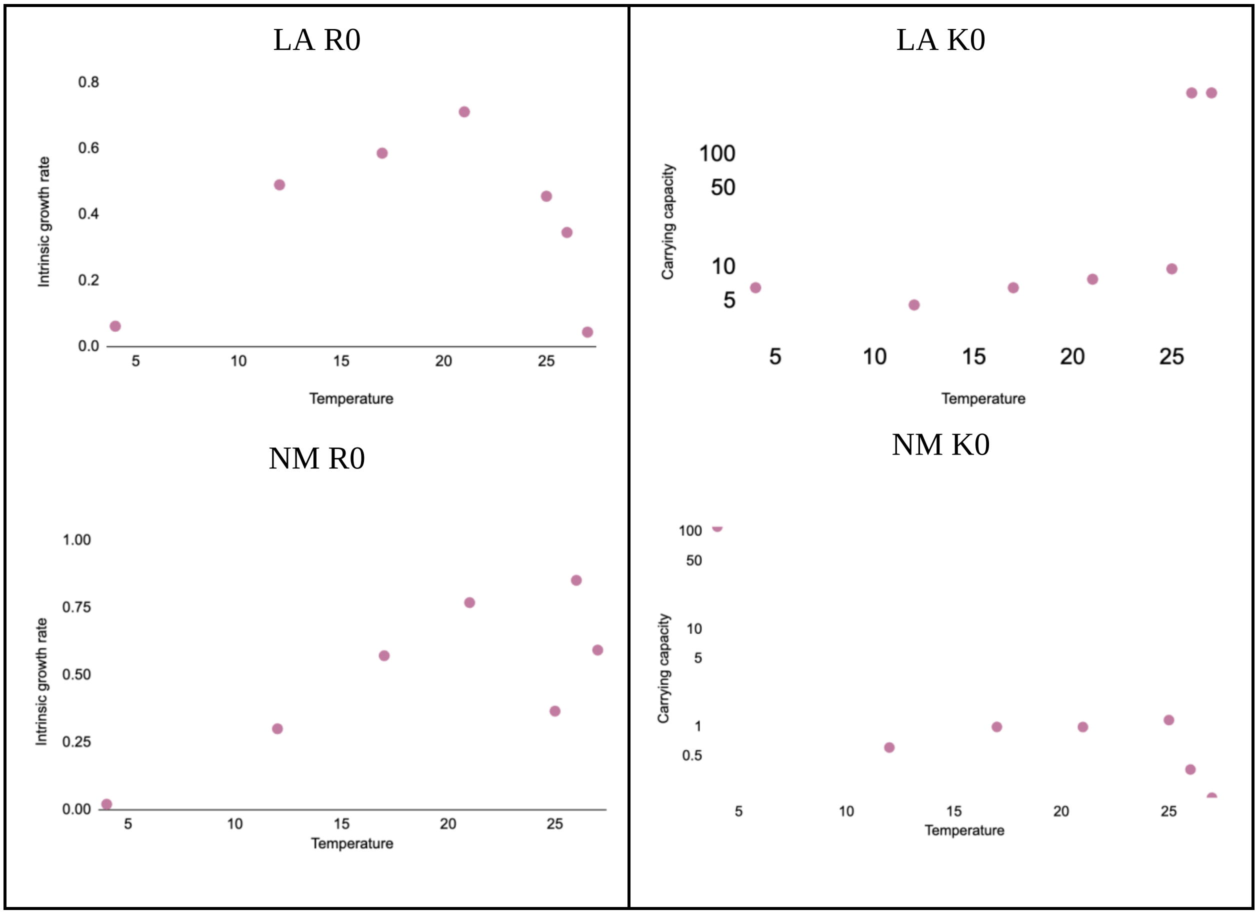

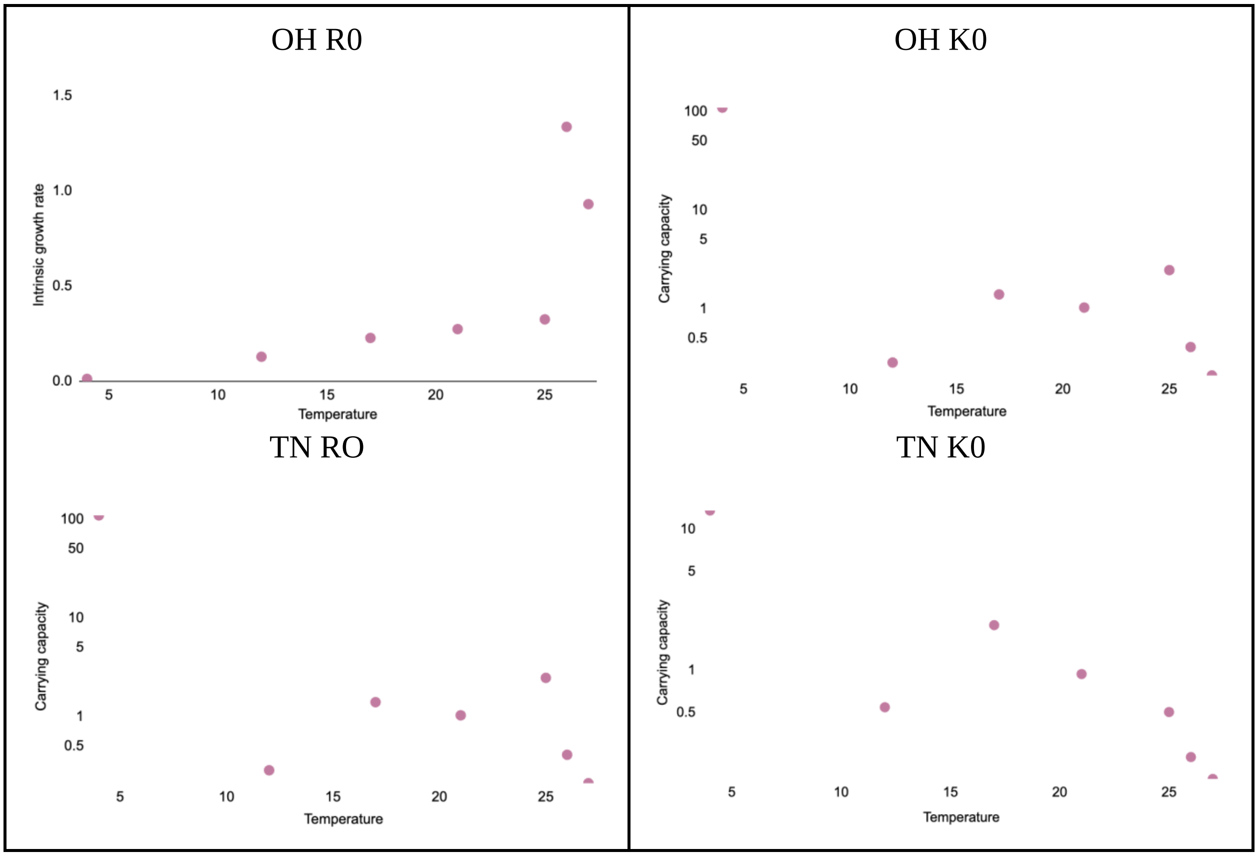


### AIC values

AIC values for models at each temperature and isolate were recorded and later averaged together. The individual AIC scores are shown here. The yellow highlights represent the model with the smallest AIC value, or the best fit

| LA isolate AIC values | | | | |
|---|---|---|---|---|
| Temp | Logistic function | Growth function | Carrying capacity function | Growth & carrying capacity function |
| 4 | -462.3636 | -458.4049 | -456.5515 | -459.3708 |
| 12 | -325.4686 | -325.9351 | -324.586 | -329.7589 |
| 17 | -130.1405 | -132.3306 | -127.8308 | -130.5935 |
| 21 | -103.1529 | -101.22 | -105.1036 | -105.2566 |
| 25 | -200.5581 | -198.1566 | -197.4213 | -195.7768 |
| 26 | -189.2776 | -200.1947 | -185.2767 | -195.8086 |
| 27 | -258.9787 | -264.156 | -254.9784 | -262.2687 |

| TN isolate AIC values | | | | |
|---|---|---|---|---|
| Temp | Logistic function | Growth function | Carrying capacity function | Growth & carrying capacity function |
| 4 | -775.0110 | -770.9966 | -771.0506 | -769.0364 |
| 12 | -411.7495 | -410.2377 | -417.2557 | -416.2248 |
| 17 | -233.4606 | -234.0631 | -231.9565 | -234.1205 |
| 21 | -189.9003 | -217.2353 | -187.8749 | -191.1274 |
| 25 | -179.9296 | -199.4041 | -190.0421 | -185.7427 |
| 26 | -204.2852 | -205.9980 | -209.5620 | -203.3215 |
| 27 | -257.2252 | -261.7935 | -253.4408 | -264.9684 |

| OH isolate AIC values | | | | |
|---|---|---|---|---|
| Temp | Logistic function | Growth function | Carrying capacity function | Growth & carrying capacity function |
| 4 | -696.6526 | -692.6929 | -692.6826 | -690.7234 |
| 12 | -475.9088 | -472.7163 | -490.6134 | -498.7264 |
| 17 | -219.8419 | -242.1374 | -222.7039 | -233.1347 |
| 21 | -187.3592 | -203.1161 | -186.1133 | -213.4574 |
| 25 | -197.5805 | -200.2775 | -194.0806 | -202.4619 |
| 26 | -171.9599 | -173.886 | -178.0197 | -173.6464 |
| 27 | -229.7017 | -231.0654 | -234.8064 | -225.4191 |

| NM isolate AIC values | | | | |
|---|---|---|---|---|
| Temp | Logistic function | Growth function | Carrying capacity function | Growth & carrying capacity function |
| 4 | -619.2097 | -615.2448 | -615.2201 | -613.2576 |
| 12 | -400.2928 | -402.5727 | -405.6032 | -399.7114 |
| 17 | -128.9328 | -128.385 | -129.0806 | -151.653 |
| 21 | -162.9391 | -178.5106 | -193.0199 | -187.8518 |
| 25 | -226.5601 | -238.3816 | -235.6971 | -230.6192 |
| 26 | -195.2315 | -210.8139 | -198.9082 | -197.6728 |
| 27 | -253.0702 | -256.6085 | -255.9606 | -256.7678 |

Since no apparent trend was recognized from the individual AIC values, they were averaged together according to isolate and temperature.

| AIC values across different isolates | | | | |
|---|---|---|---|---|
| Temp | Logistic function | Growth function | Carrying capacity function | Growth & carrying capacity function |
| 4 | -2565.2369 | -2555.3392 | -2553.5048 | -2553.3882 |
| 12 | -1625.4197 | -1629.4618 | -1656.0583 | -1665.4215 |
| 17 | -724.3758 | -754.9161 | -729.5718 | -770.5017 |
| 21 | -655.3515 | -718.082 | -690.1117 | -718.6932 |
| 25 | -816.6283 | -854.2198 | -835.2411 | -835.6006 |
| 26 | -772.7542 | -808.8926 | -789.7666 | -791.4493 |
| 27 | -1010.9758 | -1031.6234 | -1017.1862 | -1030.424 |

| AIC values across different temperatures | | | | |
|---|---|---|---|---|
| Isolate | Logistic function | Growth function | Carrying capacity Function | Growth & carrying capacity function |
| LA | -1693.9400 | -1716.398 | -1687.748 | -1720.834 |
| NM | -2010.236 | -2066.517 | -2069.49 | -2079.534 |
| OH | -2203.005 | -2251.892 | -2235.02 | -2279.569 |
| TN | -2275.561 | -2335.728 | -2297.183 | -2306.542 |

### AIC scores

To compare the varying models to the logistic function, AIC scores were calculated by averaging all 4 isolate AIC scores and subtracting their value from the logistic model, shown below.

| AIC scores across different isolates | | | |
|---|---|---|---|
| Temp | Growth function | Carrying capacity function | Growth & carrying capacity function |
| 4 | 9.8977 | 11.7321 | 11.8487 |
| 12 | -4.0421 | -30.6386 | -40.0018 |
| 17 | -30.5403 | -5.196 | -46.1259 |
| 21 | -62.7305 | -34.7602 | -63.3417 |
| 25 | -37.5915 | -18.6128 | -18.9723 |
| 26 | -36.1384 | -17.0124 | -18.6951 |
| 27 | -20.6476 | -6.2104 | -19.4482 |

| AIC scores across different temperatures | | | |
|---|---|---|---|
| Isolate | Growth function | Carrying capacity function | Growth & carrying capacity function |
| LA | -22.4580 | 6.1920 | -26.8940 |
| NM | -56.281 | -59.254 | -69.298 |
| OH | -48.887 | -32.015 | -76.564 |
| TN | -60.167 | -21.622 | -30.981 |

### AIC weights

Lastly, for the ease of presentation, the AIC scores were transformed into a probability of the model's fit relative to the acquired data and was done for the AIC score across different isolates and across different temperature ranges.

| Akaike weights across different isolates | | | |
|---|---|---|---|
| Temp | Growth function | Carrying capacity function | Growth & carrying capacity function |
| 4 | 0.007003324 | 0.002798783 | 0.00264028 |
| 12 | 1.539733E-08 | 0.009179142 | 0.9908208 |
| 17 | 0.0004125255 | 1.294214E-09 | 0.9995875 |
| 21 | 0.4241889 | 3.580023E-07 | 0.5758107 |
| 25 | 0.9998338 | 7.564069E-05 | 9.053571E-05 |
| 26 | 0.9997667 | 7.026524E-05 | 0.00016298 |
| 27 | 0.6452687 | 0.0004728708 | 0.3542372 |

| Akaike weights across different temperatures | | | |
|---|---|---|---|
| Isolate | Growth function | Carrying capacity function | Growth & carrying capacity function |
| LA | 0.09814555 | 5.896607E-08 | 0.9018531 |
| NM | 0.001478763 | 0.006538487 | 0.9919828 |
| OH | 9.772718E-07 | 2.11986E-10 | 0.999999 |
| TN | 0.9999995 | 4.266379E-09 | 4.595581E-07 |

### AIC graphs

All variations of the AIC values and scores were graphed to visually determine how well a respective model represented the data.

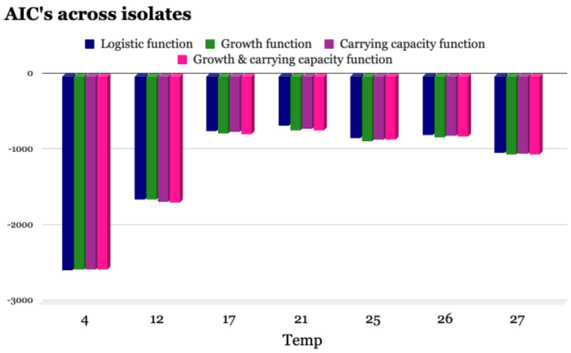


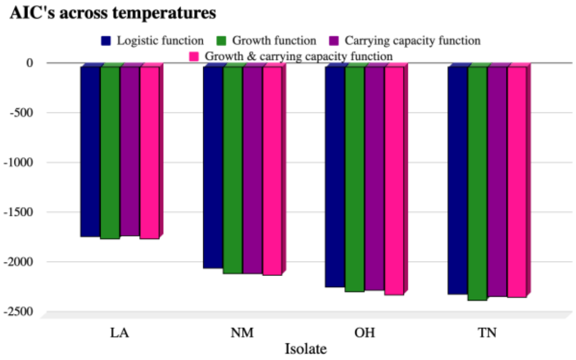


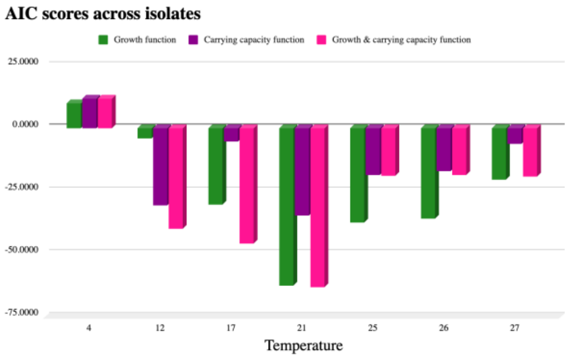


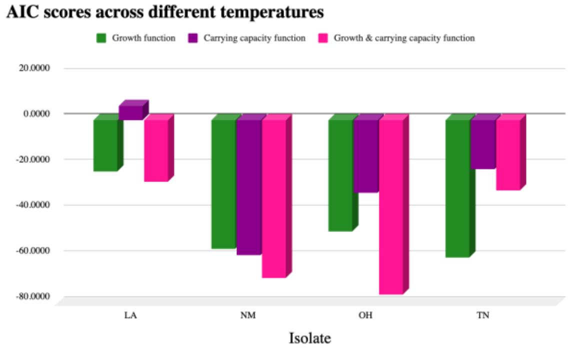
